# Interferon redundancy counteracts proteolytic inactivation by SARS-CoV-2 3CL^pro^ main protease

**DOI:** 10.64898/2026.09.07.749787

**Authors:** Peter M. Grin, Hugo C. R. de Jesus, Isabel Pablos, Reinhild Kappelhoff, Georgina S. Butler, Charaf Benarafa, Christopher M. Overall

**Affiliations:** Department of Biochemistry and Molecular Biology, University of British Columbia, Vancouver, British Columbia, V6T 1Z3, Canada; Centre for Blood Research, Life Sciences Centre, University of British Columbia, Vancouver, British Columbia, V6T 1Z3, Canada; Institute for Virology and Immunology, Mittelhäusern, 3147 Bern, Switzerland; Department of Infectious Diseases and Pathobiology, Vetsuisse Faculty, University of Bern, 3012 Bern, Switzerland; Department of Oral Biological and Medical Sciences, Faculty of Dentistry, University of British Columbia, Vancouver, British Columbia, V6T 1Z3, Canada; Multidisciplinary Center for Infectious Diseases (MCID), University of Bern, Bern, Switzerland

**Keywords:** SARS-CoV-2, COVID-19, coronavirus 3C proteases, 3CL^pro^, Mpro, interferon, glycosylation, cytokine, virus, matrix metalloproteinases, proteolysis

## Abstract

Interferons (IFNs) are secreted during virus infection and induce antiviral responses through receptor-mediated phosphorylation of signal transducer and activator of transcription (STAT) proteins, leading to IFN-stimulated gene expression with antiviral activity. We previously reported that the SARS-CoV-2 main protease, 3CL^pro^, is secreted from infected cells through gasdermin D/E pores and retains proteolytic activity in human serum against extracellular substrates. Here, we show that 3CL^pro^ selectively cleaves glycosylated IFN-λ1, IFN-λ2, and a rare naturally occurring variant of IFN-γ (Arg160Gln), but does not cleave wild-type IFN-γ, IFN-λ3, IFN-λ4, IFN-α proteins or IFN-β. We identified sites of O-linked glycosylation of IFN-λ1 at Thr^137^ and one or more threonines or serine in the sequence ^24^TSKPTTT^30^, and N-linked glycosylation at Asn^65^ that were indispensable for signaling. Unexpectedly, O-glycosylation was also required for the cleavage and inactivation of IFN-λ1 by 3CL^pro^ at two sites. Cleavage reduced STAT1 phosphorylation and impaired the induction of the IFN-stimulated proteins MX1, OAS2, and IFIT1. Although 3CL^pro^ cleaved IFN-λ2 proximal to its N-terminus at ARLH^32^↓GALP, cleavage neither disrupted signaling nor antiviral activity against SARS-CoV-2 and vesicular stomatitis virus. We further show that matrix metalloproteinases (MMPs) 2, 7, 8, and 12 degrade 3CL^pro^, whereas 3CL^pro^ shows no activity against these MMPs.

**IMPORTANCE:** Extracellular cleavage of IFN-λ1 and IFN-λ2, and the selective inactivation of IFN-λ1 by 3CL^pro^, reveals a potential pathobiological role for secreted SARS-CoV-2 3CL^pro^. The redundancy of IFN signaling, together with differential cleavage susceptibilities of IFNs to 3CL^pro^ and the degradation of 3CL^pro^ by several secreted MMPs, provide host countermeasures that preserve antiviral defence in some settings. Our findings highlight the potential to discover additional extracellular 3CL^pro^ substrates that may contribute to SARS-CoV-2 replication and COVID-19. The investigation of potential secretion, extracellular activity, and host substrates of proteases encoded by other virus families, including *Picornaviridae, Flaviviridae, Adenoviridae,* and *Poxviridae,* is also warranted. Finally, our findings support the development of cleavage-resistant IFN-λ1 as a potential broad-spectrum antiviral therapeutic.

## INTRODUCTION

Severe acute respiratory syndrome coronavirus 2 (SARS-CoV-2) replication requires proteolytic activity of the viral RNA-encoded main protease, 3C-like protease (3CL^pro^), which cleaves the viral polyprotein at 9 Leu-Gln↓ sites, one Phe-Gln↓ site, and one Val-Gln↓ site (1–4). These cleavages release functional non-structural proteins (nsp) 4 through to nsp16, enabling formation of the replication-transcription complex for expression and translation of SARS-CoV-2 structural proteins (5, 6). We and others have identified hundreds of host cell substrates of 3CL^pro^ using N-terminomics approaches (7, 8), including targets important for viral sensing and autophagy (7), and we expanded the known cleavage specificity of 3CL^pro^ to include non-canonical cleavage sites: Leu-His↓ and Leu-Met_ox_↓ (9). We recently demonstrated that the interferon (IFN)-stimulated genes (ISG) galectin-8 and 2’-5’-oligoadenylate synthetase 1 (OAS1) p46 isoform are inactivated by 3CL^pro^ cleavage (7, 10), whereas antiviral RNase L is activated by 3CL^pro^ as a failsafe “tripwire” mechanism to counteract inactivation of antiviral ISGs (10). We also showed that 3CL^pro^ is unconventionally secreted from SARS-CoV-2-infected cells through gasdermin D and gasdermin E pores, and that 3CL^pro^ remains proteolytically active in human serum despite an abundance of endogenous protease inhibitors (11). The extracellular milieu contains a multitude of potential new 3CL^pro^ substrates that remain to be investigated and that have pathobiological relevance to coronavirus disease 2019 (COVID-19).

Innate immune responses serve as the first line of defense against SARS-CoV-2 infection and are initiated by rapid pathogen recognition with induction of IFN responses to limit the spread of infection to neighbouring cells and tissues. IFNs are potent cytokines that confer antiviral protection by binding extracellularly to IFN receptors and triggering the Janus kinase (JAK)-signal transducer and activator of transcription (STAT) pathway (12, 13). STAT phosphorylation and dimerization lead to nuclear translocation, resulting in upregulation of ISG expression of hundreds of effector proteins, including interferon-induced GTP-binding protein Mx1 (MX1), OAS2, and interferon-induced protein with tetratricopeptide repeats 1 (IFIT1) (12–16). MX1 binds to viral nucleoproteins to inhibit viral replication (17, 18), whereas OAS2 detects viral RNA and activates endogenous RNase L to degrade the viral RNA (19). IFIT1 specifically binds to single-stranded RNA containing a 5’ triphosphate group to inhibit viral RNA translation (16, 20).

IFNs are classified into three types based on their cell surface receptor affinities. Type I IFNs, including thirteen IFN-α proteins and IFN-β, bind to near-ubiquitously expressed receptors composed of two distinct subunits: a low-affinity IFNAR1 that signals through tyrosine kinase 2 intracellularly, and high-affinity IFNAR2 that signals through JAK1 (21, 22). IFN-γ forms the type II IFNs that are immune cell-specific. The rare IFN-γ (Arg160Gln) variant exists naturally (SNP rs201359065) at an allele frequency ranging from T = 0.00004 – 0.0004 in East Asian and South Asian populations (dbSNP, NIH NCBI) (23). An IFN-γ dimer binds to IFNGR1 that results in ligand-induced receptor dimerization and recruitment of IFNGR2 to form the active ternary signaling complex, which signals through JAK1/2 intracellularly (24–26). Type III IFNs consist of IFN-λ1–4 that maintain epithelial barrier immunity by binding to dimeric receptors composed of IL10RB and IFNLR1, the latter of which confers binding specificity and is expressed mainly by epithelial cells of the respiratory tract, gastrointestinal tract, and oral cavity (27–29). Upon receptor binding, JAK1 activation leads to phosphorylation of STAT1 primarily at Tyr^701^ (pSTAT1-Y^701^) with downstream activation of ISGs (12, 13, 30). IFN-γ signals predominantly through phospho-(p)STAT1 homodimers that enter the nucleus and bind to γ-activated site (GAS) promoters to upregulate ISGs (31). In contrast, type I and III IFNs signal mainly through pSTAT1 and pSTAT2 heterodimers that bind to IFN regulatory factor 9 (IRF9), forming the ISGF3 complex that translocates to the nucleus and binds IFN-stimulated response elements (ISREs) to upregulate ISGs (32–34). The importance of IFN signaling in limiting SARS-CoV-2 infection is highlighted by life-threatening COVID-19 in patients with inborn errors or autoantibodies against IFNs (35, 36), and development of recombinant IFN-α-2b (37, 38) and IFN-λ (39) as antiviral drugs against COVID-19.

Matrix metalloproteinases (MMPs) are zinc-dependent proteases that regulate inflammation via proteolytic processing of many cytokines and chemokines (40–43), including IFNs (44–46). We previously showed that MMP12 plays a key role in regulating IFN-α secretion in coxsackievirus B3 infection by moonlighting as a transcription factor to upregulate NFKBIA transcription (45). MMP12 also regulates IFN-α binding to its receptor by cleaving off the C-terminal IFNAR2 binding site (45) and inactivates IFN-γ by C-terminal cleavage at Glu^135^↓Leu, which removes the IFN-γ receptor binding site and dampens JAK-STAT1 signaling (44). Furthermore, MMPs are involved in SARS-CoV-2 infection with evidence of increased MMP1, MMP2, MMP7, MMP8, and MMP14 in lung biopsies from COVID-19 patients (47). The mechanisms by which MMPs might regulate disease progression in SARS-CoV-2 infection remain incompletely understood.

In this study, we characterize 3CL^pro^ cleavage of IFN-λ1, IFN-λ2, and IFN-γ (Arg160Gln), and the functional consequences of each cleavage on IFN activity. Through detailed biochemical analysis of IFN-λ1 cleavage by 3CL^pro^, we provide evidence of three IFN-λ1 glycosylation sites and establish the effects of N-linked and O-linked glycosylation on IFN signaling and inactivating cleavage by 3CL^pro^. We also identify MMPs as host-derived extracellular antagonists of 3CL^pro^ activity via proteolytic degradation of 3CL^pro^.

## MATERIALS AND METHODS

### Recombinant protein cleavage assays

Expression and purification of recombinant 3CL^pro^ and Cys145Ala catalytic inactive mutant, each containing C-terminal 3xFLAG-Myc-6xHis tags, was performed as described in detail (7, 11). Recombinant human IFN-α-2b (STEMCELL Technologies, 78077.1), IFN-β 1a (PBL Assay Science, 11410-2), IFN-γ (Arg160Gln) variant (PeproTech, 300-02-250 µg), IFN-λ1 (R&D Systems, 1598-IL-025/CF), IFN-λ2 (R&D Systems, 1587-IL-025/CF), IFN-λ3 (R&D Systems, 5259-IL-025/CF), IFN-λ4 (R&D Systems, 9165-IF-025) and MMP12 (R&D Systems, 917-MPB-020) were obtained commercially. Upon examining the full protein sequences of the IFN-γ available from most companies, we unexpectedly found that most supply the rare IFN-γ (Arg160Gln) variant. We purchased wild-type IFN-γ from Abcam (78020). MMP1, MMP2, MMP7, MMP8, MMP9, and soluble MMP14 lacking the transmembrane anchor were expressed in CHO cells and purified in-house as described previously (48, 49). MMPs were activated by 1 mM 4-aminophenylmercuric acetate or inhibited by 50 µM marimastat for 30 min at 37°C.

IFNs or marimastat-inhibited MMPs were incubated in the presence or absence of active 3CL^pro^ (concentrations and time points listed in figure legends) in 3CL^pro^ assay buffer (150 mM NaCl, 1 mM ethylenediaminetetraacetic acid (EDTA), 2 mM dithiothreitol (DTT), 50 mM Tris, pH 7.2). Active MMPs (300 nM) were incubated with 1:10 molar ratios of 3CL^pro^ (Cys145Ala) (3 µM) in MMP Assay Buffer (150 mM NaCl, 5 mM CaCl_2_, 50 mM HEPES, pH 7.0) for 20 h at 37°C. Cleavage assays were stopped by addition of NuPAGE 1x LDS sample buffer (Thermo Fisher Scientific, NP0007) and 100 mM DTT, and samples were electrophoresed by SDS-PAGE and stained with Imperial protein stain (Thermo Fisher Scientific, 24615). Catalytically inactive 3CL^pro^ (Cys145Ala) mutant or 3CL^pro^ inhibited by 100 µM GC376 (kindly provided by Dr. John C. Vederas, University of Alberta) served as negative controls. Densitometry of scanned gels was performed using ImageJ software (version 1.52). Edman sequencing was performed at the Tufts University Protein Sequencing Core Facility to determine the N-terminal sequences of the cleavage products as previously described (50). Apparent *k*_cat_/*K*_M_ (^app^(*k*_cat_/*K*_M_) specificity constants were estimated, as previously described (51), under the assumption of a first-order reaction using the following equation:

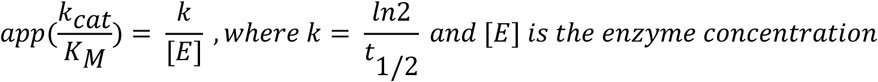

### Matrix-assisted laser desorption/ionization time-of-flight mass spectrometry

Recombinant IFN-γ (Arg160Gln) was incubated with or without 1 µM 3CL^pro^ in matrix-assisted laser desorption/ionization-time-of-flight mass spectrometry (MALDI-MS) assay buffer (8 mM Tris, 1 mM EDTA, 2 mM DTT, pH 7.0) for 20 h at 37°C before spotting onto a stainless-steel target plate, matrix (α-cyano-4-hydroxycinnaminic acid (Sigma Aldrich, 70990-1G-F)) was added and cleavage measured by MALDI-TOF MS of intact peptide substrate and cleavage products. IFN-γ peptides of the cleavage site motif, ^155^SQMLFQGRRASQ^166^YR and ^155^SQMLFRGRRASQ^166^YR, were synthesized with C-terminal Tyr-Arg residues (GenScript) to improve ionization. Peptides were incubated at 1:50 enzyme to substrate ratio with 1 µM 3CL^pro^, together with the 3CL^pro^-non-cleavable peptide NSLPYNSLPRYNSLPYNR as an internal normalization standard, for 0 – 240 min at 37°C prior to assaying by MALDI-TOF MS. All MALDI-TOF MS peptide or protein cleavage assays were performed using a 4700 Proteomics Analyzer (Applied Biosystems) in positive-ion mode and spectra were analyzed using Data Explorer Version 4.5 (Applied Biosystems).

### Liquid chromatography-tandem mass spectrometry

To identify IFN-λ1 glycosylation sites, IFN-λ1 (2.5 µg) was digested with trypsin (1:20 enzyme: substrate molar ratio). Briefly, tryptic peptides were desalted using Oasis HLB cartridges (Waters, 186000383) and 1 µg was directly loaded onto a C18 Aurora column (25 cm x 50 µm ID, 1.6 µm, IonOpticks) and eluted on an Easy nLC-1000 (Thermo Fisher Scientific) as previously described (11). Eluted peptides were ionized on an Impact II Ultra-High Resolution Q-TOF instrument (Bruker Daltonics) and data-dependent acquisition was used to obtain MS2 spectra that were analyzed with Byonic software (Protein Metrics, Version 4.4).

### Cell lines and endogenous IFN signaling

THP-1 human monocytic cells (ATCC, TIB-202) were maintained in RPMI-1640 Medium (Sigma, R8758) supplemented with 10% v/v fetal bovine serum (FBS; Sigma, F1051) and 1% v/v Penicillin-Streptomycin (Pen-Strep; Gibco, 15140122). HEK 293T cells (ATCC, CRL-3216) were cultured in Dulbecco’s modified Eagle’s medium (DMEM) with GlutaMAX and 25 mM HEPES (Gibco, 32430-100), supplemented with 10% FBS (Serana, S-FBS-MX-015) and 1% Pen-Strep. HEK-Blue™ IFN-λ Reporter Cells (InvivoGen, hkb-ifnlv2) were cultured in the same media as HEK 293T cells, with antibiotic selection using 1 µg/mL puromycin (InvivoGen, ant-pr), 10 µg/mL blasticidin (InvivoGen, ant-bl), and 100 µg/mL zeocin (InvivoGen, ant-zn). Caco-2 human intestinal epithelial cells (ATCC, HTB-37) were maintained in DMEM (Sigma, D6429) with 10% FBS, 1% Pen-Strep, and 1x GlutaMAX (Gibco, 35050-061) for STAT1 signaling experiments, and in Minimum Essential Medium (MEM) Eagle (PAN Biotech, P04-08500) with 10% FBS (Serana, S-FBS-MX-015) and 1% Pen-Strep for VSV infection experiments. A549-hACE2 cells were maintained in DMEM with GlutaMAX and 25 mM HEPES (Gibco, 32430-100), supplemented with 10% FBS (Serana, S-FBS-MX-015), 1% Pen-Strep, 1x MEM non-essential amino acids (NEAA; Gibco, 11140-035), and 1 mM sodium pyruvate (Gibco, 11360-039), with antibiotic selection using 1 µg/mL puromycin. Vero E6 cells expressing transmembrane serine protease 2 (TMPRSS2) (NIBSC Research Reagent Repository, 100978) were maintained in DMEM with GlutaMax and 25 mM HEPES, supplemented with 10% FBS (Serana, S-FBS-MX-015), 1x NEAA, and 1% Pen-Strep, with antibiotic selection using 200 µg/mL geneticin (G418 sulfate; Gibco, 10131027). Calu-3 human lung epithelial cells (ATCC, HTB-55) were maintained in MEM Alpha (Gibco,12571-063) supplemented with 20% FBS (Sigma, F1051), 1% Pen-Strep, 1x GlutaMAX, 1x Antibiotic-Antimycotic (Gibco, 15240-062). HepG2 human liver cells (ATCC, HB-8065) were maintained in DMEM with 10% FBS and 1% Pen-Strep. All cell lines used in this study were cultured at 37°C with 5% CO_2_ and were passaged using 0.25% trypsin (Gibco, 15090-046) or TrypLE Express for Calu-3 cells (Gibco, 12605-28), and tested negative for mycoplasma contamination.

THP-1 cells were differentiated into M0-like macrophages with 100 ng/mL phorbol 12-myristate 13-acetate for 24 h, washed 3x with phosphate-buffered saline (PBS), then serum-starved and treated with 0.01–20 ng/mL of 3CL^pro^-cleaved or intact IFN-γ in the presence of polymyxin B (10 µg/mL; Cayman Chemical, 14157-500) to remove any lipopolysaccaride. Caco-2, Calu-3, or HepG2 cells were serum-starved and then treated with 3CL^pro^-cleaved or intact IFN-λ1 or IFN-λ2 (1 µg/mL) in the presence of polymyxin B (10 µg/mL) for the indicated time points. Cells were lysed on ice with 1x RIPA buffer (Abcam, ab156034) containing 1x protease inhibitor cocktail (Bimake.com, B14001), 1x phosphatase inhibitor cocktail A (Bimake.com, B15001-A), and 1x phosphatase inhibitor cocktail B (Bimake.com, B15001-B), then 1x LDS sample buffer and 100 mM DTT were added to prepare samples for gel electrophoresis.

### SDS-PAGE and immunoblotting

Lysates were sonicated for 1–3 s using a Sonic Dismembrator Model 100 at intensity 3 (Fisher Scientific) and heated at 70°C for 5 – 10 min, then electrophoresed on Bolt 8% Bis-Tris Plus gels (Thermo Fisher Scientific, NW00080), or 12% NuPAGE Bis-Tris gels (Thermo Fisher Scientific, NP0341) at 200 V for 35 – 50 min in 1x MES SDS Running Buffer (50 mM MES, 50 mM Tris base, 0.1% SDS, 1 mM EDTA, pH 7.3). Separated proteins were transferred to methanol-activated PVDF membrane (Immobilon-FL, Millipore, IPFL00010) in 1x NuPAGE Transfer Buffer (25 mM Bicine, 1 mM EDTA, 25 mM Bis-Tris, pH 7.2) at 20 V for 1 h. Membranes were blocked with Odyssey Intercept Protein-Free Blocking buffer (Li-COR Biosciences, 927-90001) for 1 h at room temperature, then incubated with primary antibodies (see Table 1) diluted in Odyssey Intercept buffer with 0.2% Tween-20 overnight at 4°C (or at room temperature for 2 h). Blots were washed 3x with Tris-buffered saline containing 0.1% Tween-20 (TBS-T), incubated for 1 h at room temperature with secondary antibodies (Table 1) coupled to AlexaFluor 680 (Thermo Fisher Scientific) or IR-Dye 800 (Li-COR Biosciences) and diluted 1:10,000 in Odyssey Intercept buffer with 0.2% Tween-20 and 0.01% SDS, washed an additional 3x in TBS-T then scanned using an Odyssey Infrared Imager (Li-COR Biosciences). Densitometry of immunoreactive bands was performed using Image Studio Lite software (Li-COR Biosciences).

**Table 1.** List of antibodies used for immunoblotting for various target proteins.

| Target Protein | Host Species | Dilution | Type/Clonality (Clone ID) | Source (Catalogue #; RRID) |
| --- | --- | --- | --- | --- |
| IFN-λ1-3 | Goat | 1:500 | Polyclonal | (AF1598; RRID:AB_354883) |
| pSTAT1-Y<sup>701</sup> | Rabbit | 1:1000 | Monoclonal (58D6) | Cell Signaling (9167) |
| STAT1 | Mouse | 1:1000 | Monoclonal (SM1) | Abcam (ab3987; RRID:AB_304210) |
| β-actin | Mouse | 1:2000 | Monoclonal | Abcam (ab8226; RRID:AB_306371) |
| β-tubulin | Mouse | 1:2000 | Monoclonal (BT7R) | UBC Antibody Lab |
| MX1 | Rabbit | 1:500 | Polyclonal | Proteintech (13750-1-AP; RRID:AB_2266768) |
| OAS2 | Rabbit | 1:500 | Polyclonal | Proteintech (19279-1-AP; RRID:AB_10642832) |
| IFIT1 | Rabbit | 1:500 | Polyclonal | Proteintech (23247-1-AP; RRID:AB_2811269) |
| iNOS | Rabbit | 1:500 | Polyclonal | Proteintech (18985-1-AP; RRID:AB_2782960) |
| RSAD2 | Rabbit | 1:500 | Polyclonal | ABclonal (A8271) |
| BST2 | Rabbit | 1:500 | Polyclonal | Proteintech (13560-1-AP; RRID:AB_2067220) |
| SOCS1 | Rabbit | 1:500 | Polyclonal | ABclonal (A7754) |
| Rabbit IgG (H+L) AlexaFluor 680 | Goat | 1:10,000 | Secondary | Invitrogen (A-21109; RRID:AB_2535758) |
| Mouse IgG<br>IRDye 800CW | Donkey | 1:10,000 | Secondary | Li-COR Biosciences<br>(926-32212; RRID:AB_621847) |

## Antibodies

### Transfection and HEK-Blue IFN-**λ** Signaling

HEK 293T cells were seeded in T25 flasks overnight and grown to ∼70–80% confluence. Transfections were performed in OptiMEM I Media (Gibco, 31985-062) using TurboFect Transfection Reagent (ThermoFisher Scientific, R0531) and 3.75 µg of IFN**-**λ2 plasmid DNA (GenScript, clone OHu17422) or IFN-λ2 Δ1-7 cleaved analogue that was generated in the same plasmid vector (GenScript) and confirmed by Sanger sequencing using T7 forward (5’-TAA TAC GAC TCA CTA TAG G-3’) and BGH reverse (5’-TAG AAG GCA CAG TCG AGG-3’) primers (Microsynth). After 48-h transfection, the conditioned media were harvested and centrifuged at 350 x g for 10 min to pellet any lifted cells, and then aliquoted and stored frozen at -70°C. IFN-λ2 concentrations were quantified by ELISA (R&D Systems, DY1598B) and confirmed by Western blot (R&D Systems, anti-IFN-λ1 polyclonal antibody, 1:500). HEK-Blue™ IFN-λ Reporter Cells (Invitrogen, hkb-ifnlv2) were treated with conditioned media containing equivalent concentrations of IFN-λ2 or Δ1-7 cleaved analogue as described in figure legends; time-course experiments involved overnight induction of reporter gene expression following designated treatment time with IFNs as indicated. Conditioned media from IFN-stimulated HEK-Blue cells were diluted 1:10 in Quanti-Blue reagent (InvivoGen, rep-qbs) and incubated at 37°C for 1 h, and then the absorbance was measured at 650 nm using a Tecan Infinite F50 plate reader.

### Virus infections

Confluent A549-hACE2 cells were treated for 24 h with 100 or 1000 ng/mL of IFN-λ2 or Δ1-7 cleaved analogue at 37°C in serum-free and selection-free maintenance media, then inoculated with a multiplicity of infection (MOI) of 1.0 of SARS-CoV-2 D614G for 1 h at 37°C in a Biosafety Level 3 laboratory in compliance with approved biosafety protocols at the Institute for Virology and Immunology, Switzerland. The cells were washed 3x with warm PBS to remove any residual virus. Viral replication was quantified in the conditioned media at 48 h post-infection by endpoint dilution assay on TMPRSS2-expressing Vero E6 cells as described previously (52), by incubating these cells with serial 10-fold dilutions of conditioned media for 3 days at 37°C. Conditioned medium was then aspirated, and cells were fixed for at least 10 min with 4% v/v neutral-buffered formalin and stained with crystal violet. Infected wells were counted based on evidence of cytopathic effects (cell detachment), and the TCID_50_/mL was calculated using the standard Spearman-Karber method.

Caco-2 cells were grown to 80-90% confluence in 96-well plates and were treated with 1 – 1000 ng/mL of IFN-λ2 or Δ1-7 cleaved analogue for 24 h at 37°C in serum-free maintenance media. Conditioned media were aspirated and the cells were infected with a MOI of 0.5 of replication-deficient (ΔG) VSV engineered to express firefly luciferase (53) for 24 h. Cells were lysed for 10 min on ice with 1x Firefly Luciferase Lysis Buffer (Biotium, 99923), then mixed at 1:1 (v/v) ratio with 5-fold diluted ONE-Glo Luciferase Assay substrate (Promega, E6120) in opaque, white 96-well plates. Luminescence was measured using a Promega GloMax plate reader at 1 s intervals between wells.

### Enzymatic deglycosylation

Recombinant IFN-λ1 was incubated at 37°C with PNGase F (500 units/reaction, New England Biolabs, P0704S), O-glycosidase (120,000 units/reaction, New England Biolabs, E0540S), α2-3,6,8 neuraminidase (100 units/reaction, New England Biolabs, E0540S) or combinations thereof for 20 or 24 h (specified in figure legends). In 3CL^pro^ cleavage experiments, native or de-glycosylated IFN-λ1 reaction mixtures were incubated in the presence or absence of 5 µM 3CL^pro^ for 20 h at 37°C prior to SDS-PAGE analysis.

### Protein modeling

AlphaFold predictions of the IFN-λ1 (model ID: AF-Q8IU54-F1) and IFN-λ2 (model ID: AF-Q8IZJ0-F1) structures were modeled using PyMOL (Schrodinger LLC, version 2.3.4) to illustrate the disordered N-termini that were not resolved in reported structures of IFN-λs (54, 55). Schematics showing the effects of IFN-λ1 glycosylation on cleavage by 3CL^pro^, and the models of MMP inactivation of 3CL^pro^ and IFN-λ2 activation by 3CL^pro^, were made using BioRender.com.

### Statistical analyses

Data were plotted as mean ± SD and analysed by one-way ANOVA with Tukey’s post-hoc test for multiple comparisons, with a significance threshold of p <0.05 (GraphPad Prism, version 10.1.0). Independent biological replicates are denoted by *n* and consist of independently treated wells for cell experiments or independent co-incubations for biochemical experiments. Independent experiments are denoted by *N* and represent experiments conducted under similar conditions on different days. All *N* and *n* values are reported in the figure legends.

## RESULTS

### 3CLpro cleaves the IFN-**γ** (Arg160Gln) variant

We determined whether type I or II IFNs are cleaved and inactivated by SARS-CoV-2 3CL^pro^ as a potential mechanism for dampening antiviral host defences. 3CL^pro^ was incubated with recombinant human IFN-α-2b, IFN-β, and IFN-γ (wild-type, WT) or the IFN-γ (Arg160Gln) variant. IFN-α-2b was selected from the 13 human IFN-α proteins because it contains more consensus 3CL^pro^ cleavage sites (LQ/LH/LM_ox_↓) (9) than other IFN-αs, and is also in development as an antiviral drug for treating and prophylaxis against COVID-19 (37, 38).

SDS-PAGE analyses of cleavage assays revealed that IFN-γ (Arg160Gln) was susceptible to cleavage by 3CL^pro^, whereas IFN-α-2b, IFN-β, and wild-type IFN-γ were not (Fig. 1a). 3CL^pro^ progressively cleaved IFN-γ (Arg160Gln) over time as evidenced by a decrease in full-length IFN-γ (Arg160Gln) on 12% SDS-PAGE (Fig. 1b). This reduction occurred with the concomitant appearance of an ∼18-kDa band that, by Edman degradation, shared the same N-terminal sequence as full-length IFN-γ (Arg160Gln) (Fig. 1a), thereby indicating a C-terminal cleavage event by 3CL^pro^. The negative controls, catalytically inactive 3CL^pro^ (Cys145Ala) mutant and 3CL^pro^ inhibited by GC376, showed no cleavage (Fig. 1b) (56). Densitometric quantification and kinetic analyses of the cleavage products showed that 3CL^pro^ cleaved IFN-γ (Arg160Gln) with an ^app^(*k*_cat_/*K*_M_) of 3.33 x 10^2^ M^-1^s^-1^ (Fig. 1c). Top-down MALDI-TOF MS analysis of the 3CL^pro^-cleaved IFN-γ Arg160Gln identified a cleaved peptide product with a *m/z* of 674.5 (Fig. 1d). This peptide accurately corresponds to the molecular weight (± 0.2 *m/z*) of the C-terminal sequence, ^161^GRRASQ-COOH, which is the predicted product generated by proteolysis at MLFQ^160^↓GRRA.

**FIG 1.**
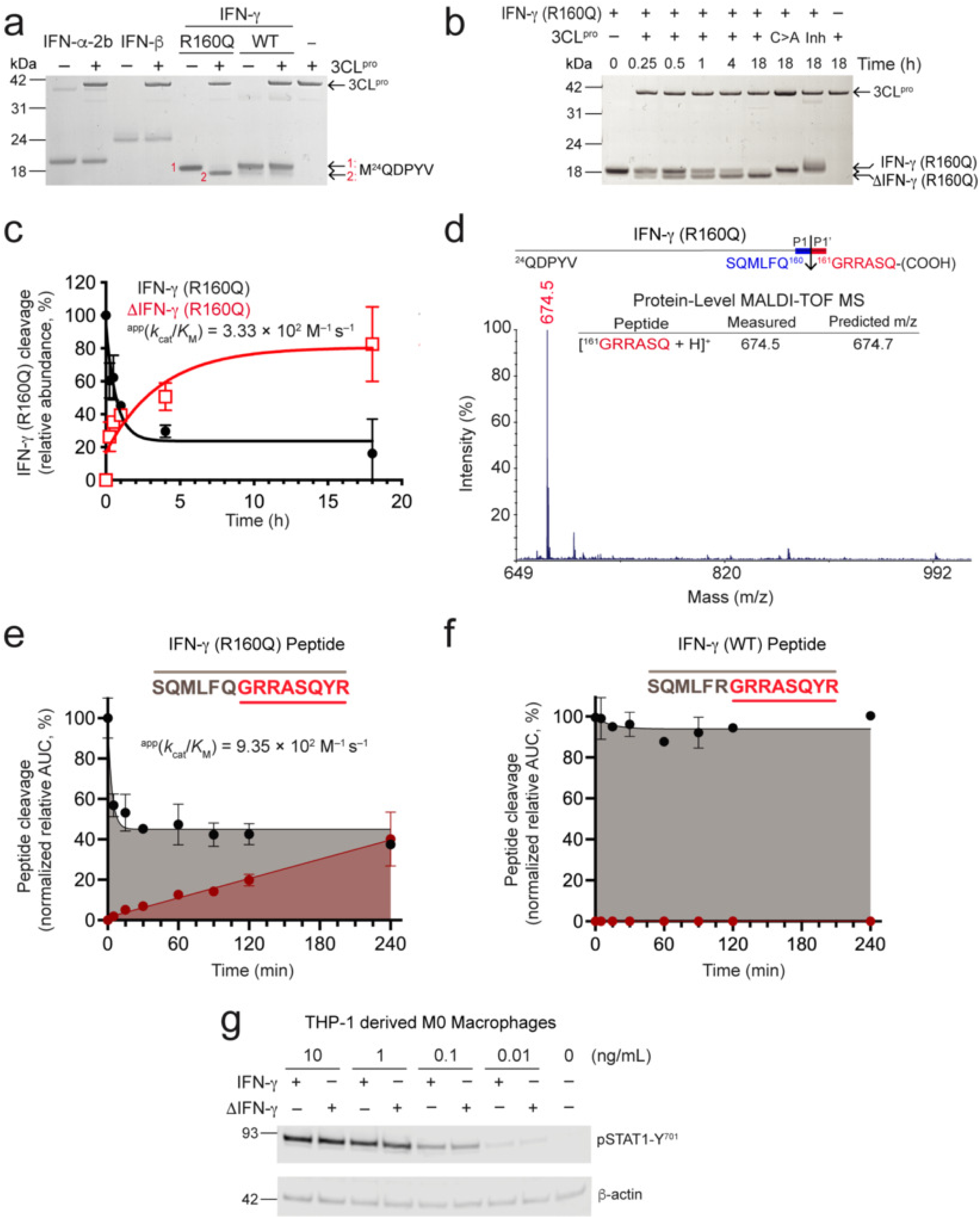
SARS-CoV-2 3CL^pro^ efficiently cleaves the IFN-γ (Arg160Gln) variant but not IFN-α-2b or IFN-β. (a) *In vitro* incubation of recombinant human IFN-α-2b, IFN-β, and wild-type (WT) IFN-γ or the IFN-γ (Arg160Gln) variant with recombinant 3CL^pro^ (1 µM) or assay buffer for 18 h at 37°C, and analysed by Coomassie-stained 12% SDS-PAGE (*N* = 2). N-terminal sequences of intact (1) or cleaved (2) IFN-γ (Arg160Gln) were identified by Edman microsequencing, M^24^, initiator Met. *, non-specific protein band (b) Time-course cleavage of IFN-γ (Arg160Gln) by 3CL^pro^ (1 µM) over 18 h at 37°C. C>A, catalytically inactive 3CL^pro^ (Cys145Ala); Inh, 3CL^pro^ inhibited by 100 µM GC376. Reaction products were resolved by 12% SDS-PAGE and stained with Coomassie R-250 (*N* = 2). (c) Densitometric analysis of IFN-γ (Arg160Gln) cleavage over time showing a decrease in full-length IFN-γ (Arg160Gln) (black, filled circles) accompanied by a concomitant increase in ΔIFN-γ (Arg160Gln) (red, open squares). The calculated apparent specificity constant (^app^(*k*_cat_/*K*_M_)) for 3CL^pro^ is shown. Data are plotted as mean ± range (*n* = 2, *N* = 2). (d) Protein-level MALDI-TOF MS spectra of 3CL^pro^ cleavage assay products of IFN-γ (Arg160Gln). The cleaved C-terminal peptide, ^161^GRRASQ, at the C-terminus with a m/z 674.5 was identified. Schematic diagram of the mature N-terminal, P6–P6’, and non-prime (P6 – 1, blue) and prime side (P1’ – P6’, red) sequences of the 3CL^pro^ cleavage site, indicated by i, with the IFN-γ C-terminal peptide cleavage product and its corresponding m/z predicted and measured are shown. (e and f) Enzyme kinetic parameters ^app^(*k*_cat_/*K*_M_) of 3CL^pro^ IFN-γ peptide cleavage by MALDI-TOF measurements of P′–product generation (red) and substrate consumption (black). (e) P6 – P6’ of IFN-γ (Arg160Gln) and (f) IFN-γ wild-type sequences, with C-terminal Tyr-Arg adapters used to improve ionization (*n* = 2, *N* = 2). (g) Immunoblots for pSTAT1-Y^701^ in cell lysates collected from THP-1-derived macrophages treated with intact IFN-γ (Arg160Gln) or 3CL^pro^-cleaved IFN-γ (Arg160Gln) (ΔIFN-γ) for 1 h at 37°C (*N* = 2). β-actin loading control and molecular weight markers in kDa are shown. Immunoblot antibodies and dilutions used: anti-pSTAT1-Y^701^ (1:1000), anti-β-actin (1:2000). See Fig S6 for full uncropped gels and immunoblots.

Cleavage of a synthetic peptide, ^155^SQMLFQGRRASQ^166^YR, designed from this sequence up to and including the carboxyl-terminal amino acid residue (Gln^166^), confirmed the cleavage site at P1-Gln^160^. Kinetic analysis of time-course peptide cleavage assays estimated the ^app^(*k*_cat_/*K*_M_) to be 9.35 × 10^2^ M^-1^s^-1^ (Fig. 1e), in agreement with cleavage kinetics of the intact protein cleavage and ranking it among the most efficiently cleaved of > 200 sites measured (7, 9), including RPAP1, IRS2, PTBP1 (7), gasdermin D (11), and OAS1-p46 (10). In contrast, cleavage did not occur in the control peptide of WT IFN-γ with Arg^160^ (underlined), ^155^SQMLF<u>R</u>GRRASQ^166^YR (Fig. 1f). These data show that the variant IFN-γ (Arg160Gln) is an efficiently cleaved 3CL^pro^ substrate, whereas IFN-α, IFN-β, and WT IFN-γ are resistant.

### 3CLpro does not inactivate IFN-**γ**

To determine whether 3CL^pro^-mediated cleavage of IFN-γ (Arg160Gln) alters STAT1 signaling during M1-like macrophage polarization, we first differentiated THP-1 monocytes into M0 macrophage-like cells using phorbol 12-myristate 13-acetate and then treated the cells to promote M1-like polarization by IFN-γ. Both 3CL^pro^-cleaved and intact IFN-γ (Arg160Gln) induced comparable STAT1 phosphorylation en route to M1 polarization across a broad range of times from 5 min to 24 h post-treatment (Fig. 1g and S1a–f). These results indicate that removal of the six C-terminal residues does not impair IFN-γ-mediated STAT1 signaling.

### 3CLpro cleaves IFN-**λ**1 and IFN-**λ**2, but not IFN-**λ**3

The IFN-λ family of cytokines, consisting of IFN-λ1–4, plays important antiviral roles by promoting epithelial barrier function at mucosal surfaces in the respiratory and gastrointestinal tracts (57). Previously we identified two 3CL^pro^ cleavage sites in IFN-λ1 at SQLQ^129^↓ACIQ and HRLQ^154^↓EAPK (Fig. S2) using amino terminal-oriented mass spectrometry (11). Therefore, we analyzed all human IFN-λs for cleavage susceptibility. Incubation of full-length recombinant IFN-λ1 (20–200) with increasing concentrations of 3CL^pro^ generated increased amounts of an ∼15-kDa cleavage product consistent with IFN-λ1 (1–129) (Fig. 2a) (11). The predicted MW of mature IFN-λ1 is 20,018 Da, but with an apparent electrophoretic mobility of the protein on 12% SDS-PAGE ranging from ∼26 to ∼35 kDa this indicated significant post-translational modification of the translated protein (Fig. 2a). 3CL^pro^ had a reduced effect on cleavage of these proteoforms, suggesting that post-translational modifications (PTMs) of IFN-λ1 alter cleavage susceptibility.

**FIG 2.**
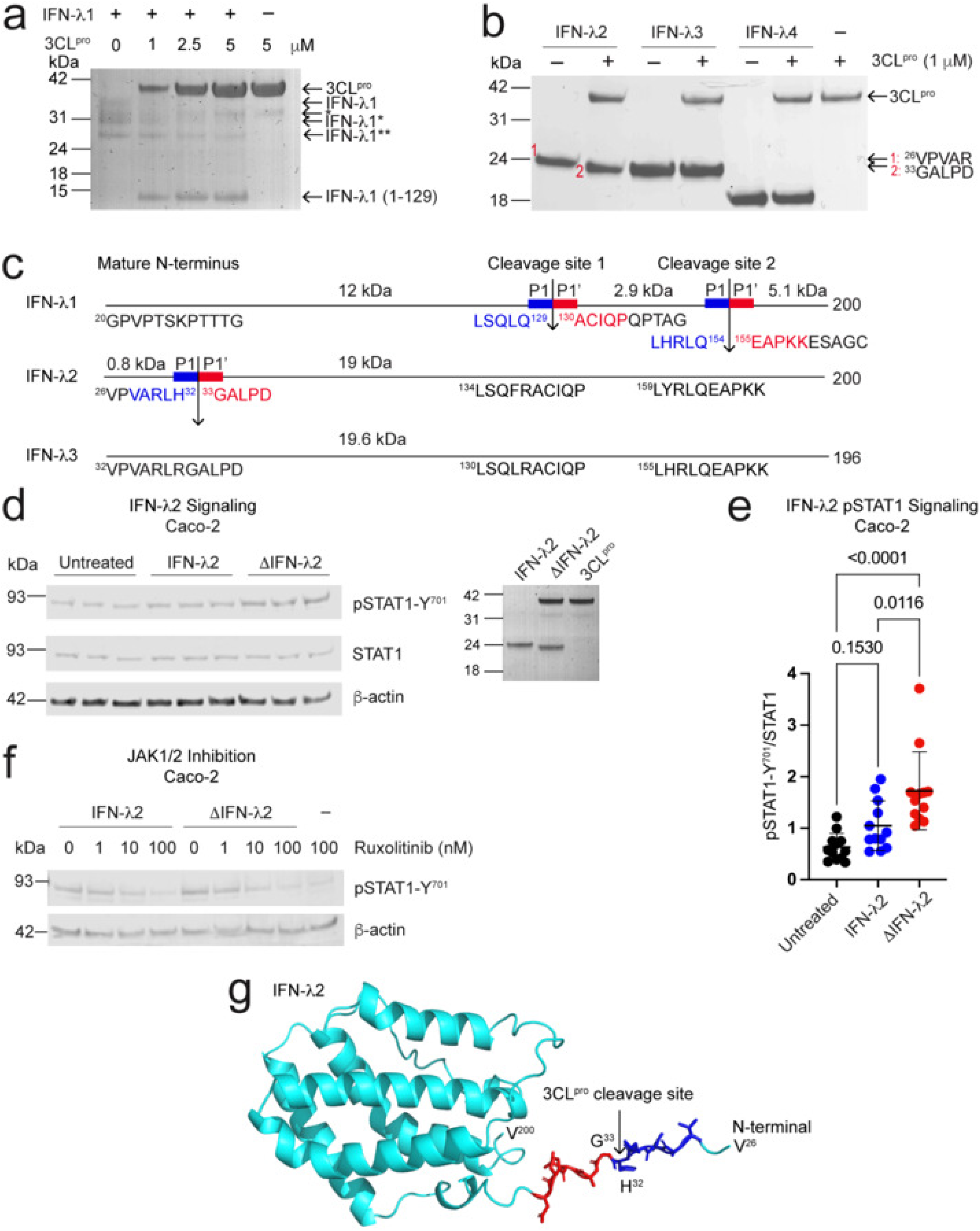
SARS-CoV-2 3CL^pro^ cleaves IFN-λ1 and IFN-λ2, but not IFN-λ3 and IFN-λ4. (a) 12% SDS-PAGE separation of recombinant human IFN-λ1 (1 µg) incubated with increasing concentrations of active 3CL^pro^ for 18 h at 37°C (representative of *N* = 2 experiments). IFN-λ1 (1–129), 3CL^pro^ cleavage product. IFN-λ1**, lowest molecular weight form indicating different post-translational modifications, including glycosylation, compared with higher MW forms denoted as IFN-λ1* or IFN-λ1, which electrophoresed with an apparent MW of ∼29 or 35 kDa, respectively, versus predicted MW of 20,018 Da; *, non-specific protein band. (b) *In vitro* incubation of recombinant human IFN-λ2, IFN-λ3, or IFN-λ4 with or without 3CL^pro^ (1 µM) for 20 h at 37°C, analyzed by 12% SDS-PAGE (*N* = 2). Numbered bands, N-terminal sequences of intact (1) or cleaved (2) IFN-λ2 were identified by Edman microsequencing, showing the mature N-terminal sequence of IFN-λ2 (^26^VPVAR; band #1 in red) and a neo-N-terminal sequence for cleaved IFN-λ2 (^33^GALPD; band #2 in red). (c) Schematic illustrations of human IFN-λ1, IFN-λ2, or IFN-λ3 showing corresponding sequences of their mature N-terminus, P5–P1, and P1’–P5’ of 3CL^pro^ cleavage sites (blue and red, respectively). The LSQ and RLQ motifs are conserved in IFN-λ1, IFN-λ2, or IFN-λ3, but substitution of unfavored P1’ residues corresponding to site 1 in IFN-λ2 and IFN-λ3 precluded cleavage here and subsequent cleavage at site 2. (d) Immunoblots for pSTAT1-Y^701^, STAT1, and β-actin in lysates prepared from Caco-2 cells treated with 1 µg/mL of IFN-λ2 or 3CL^pro^-cleaved IFN-λ2 (ΔIFN-λ2) for 1 h at 37°C (left panel; representative of *n* = 12 biological replicates per group from *N* = 4 independent experiments). 12% SDS-PAGE gel showing complete cleavage of IFN-λ2 used in the cell experiment (right panel). (e) Densitometric quantification of pSTAT1-Y^701^ relative to STAT1 in Caco-2 lysates (mean ± SD; *n* = 12, *N* = 4) with statistical significance assessed using one-way ANOVA with Tukey’s post-hoc test (p < 0.05 significance threshold). (f) Caco-2 cells were pre-treated for 2 h at 37°C with 0 – 100 nM concentrations of JAK1/2 inhibitor ruxolitinib (Rux), then treated for 1 h with 1 µg/mL of intact IFN-λ2 or ΔIFN-λ2, and lysates were immunoblotted for pSTAT1-Y^701^ and β-actin (*N* = 2). (g) AlphaFold structure prediction of IFN-λ2 (cyan) showing N-terminal 3CL^pro^ cleavage site with non-prime side in blue and prime side in red (model ID: AF-Q8IZJ0-F1). For all blots and SDS-PAGE gels, loading control and molecular weight markers in kDa are shown. Immunoblot antibodies and dilutions used: anti-pSTAT1-Y^701^ (1:1000),anti-STAT1 (1:1000), anti-β-actin (1:2000). See Fig. S7 for full uncropped gels and immunoblots.

IFN-λ2, IFN-λ3, or IFN-λ4 were also incubated with or without 3CL^pro^. SDS-PAGE and Edman sequencing identified a single cleavage of IFN-λ2 at ARLH^32^↓GALP, resulting in the removal of a 7-residue N-terminal peptide (Fig. 2b, c). Histidine substitutes for P1-Gln in ∼10% of previously identified 3CL^pro^ cleavage sites. P1-Arg is not a scissile residue (7, 9), explaining the absence of cleavage at the homologous sequence in IFN-λ3. Homologous sequences of site 1 in IFN-λ1 occur in IFN-λ2 and IFN-λ3, but with an Arg^138^ in P1 (Fig. 2c, S2), thereby conferring resistance to cleavage here. At the identical motif 2, IFN-λ2 and IFN-λ3 are not cleaved, whereas in IFN-λ1, this noncanonical site is slowly cut (Fig. 2b). These findings support a model in which cleavage at site 1 of IFN-λ1 destabilizes its structure and exposes site 2, which is otherwise occluded within the native fold. Because site 1 is not cleaved in IFN-λ2 or IFN-λ3, their cleavable site 2 remains inaccessible. Finally, IFN-λ4, with a very low homology to IFN-λs 1 – 3 of between 28 and 29 % (Fig. S2), does not contain any 3CL^pro^ cleavage motifs, and was not cleaved. The resistance of IFN-λ3 and IFN-λ4 further supports the substrate specificity of 3CL^pro^ for IFN-λ1 and IFN-λ2.

### IFN-λ2 cleavage by 3CL^pro^ does not impact antiviral signaling

To determine whether 3CL^pro^ cleavage of IFN-λ2 modulates signaling, we treated Caco-2 (colon adenocarcinoma) epithelial cells with IFN-λ2 and compared STAT1 phosphorylation at Tyr^701^ (pSTAT1-Y^701^) with 3CL^pro^-cleaved IFN-λ2. Unexpectedly, pSTAT1-Y^701^ increased relative to total STAT1 in cells treated with 3CL^pro^-cleaved IFN-λ2 (*n* = 12, *N* = 4 independent experiments) (Fig. 2d, e). Nonetheless, the magnitude of response was modest relative to basal STAT1 phosphorylation in positive controls. JAK1/2 inhibition by ruxolitinib reduced pSTAT1-Y^701^ signaling in a dose-dependent manner for cells treated with intact or cleaved IFN-λ2 (Fig. 2f), confirming signal activity via this pathway. These results suggest that removal of the unstructured N-terminal 7 amino acids from IFN-λ2 (Fig. 2g) has only a modest effect on STAT1 phosphorylation.

We next expressed the cleaved analogue of IFN-λ2 lacking the N-terminal 7 residues (Val26 to His32, hereafter referred to as ΔIFN-λ2) in HEK 293T cells. The conditioned media was used to stimulate HEK-Blue IFN-λ reporter cells. These cells express the IFNLR1 and IL10RB receptor subunits and the signaling proteins needed to activate the ISRE promoter, which drives secretion of alkaline phosphatase as the reporter enzyme. The 3CL^pro^-cleaved analogue (ΔIFN-λ2), induced signaling similar to that of full-length IFN-λ2 across a broad range of concentrations and treatment times (Fig. 3a, b).

**FIG 3.**
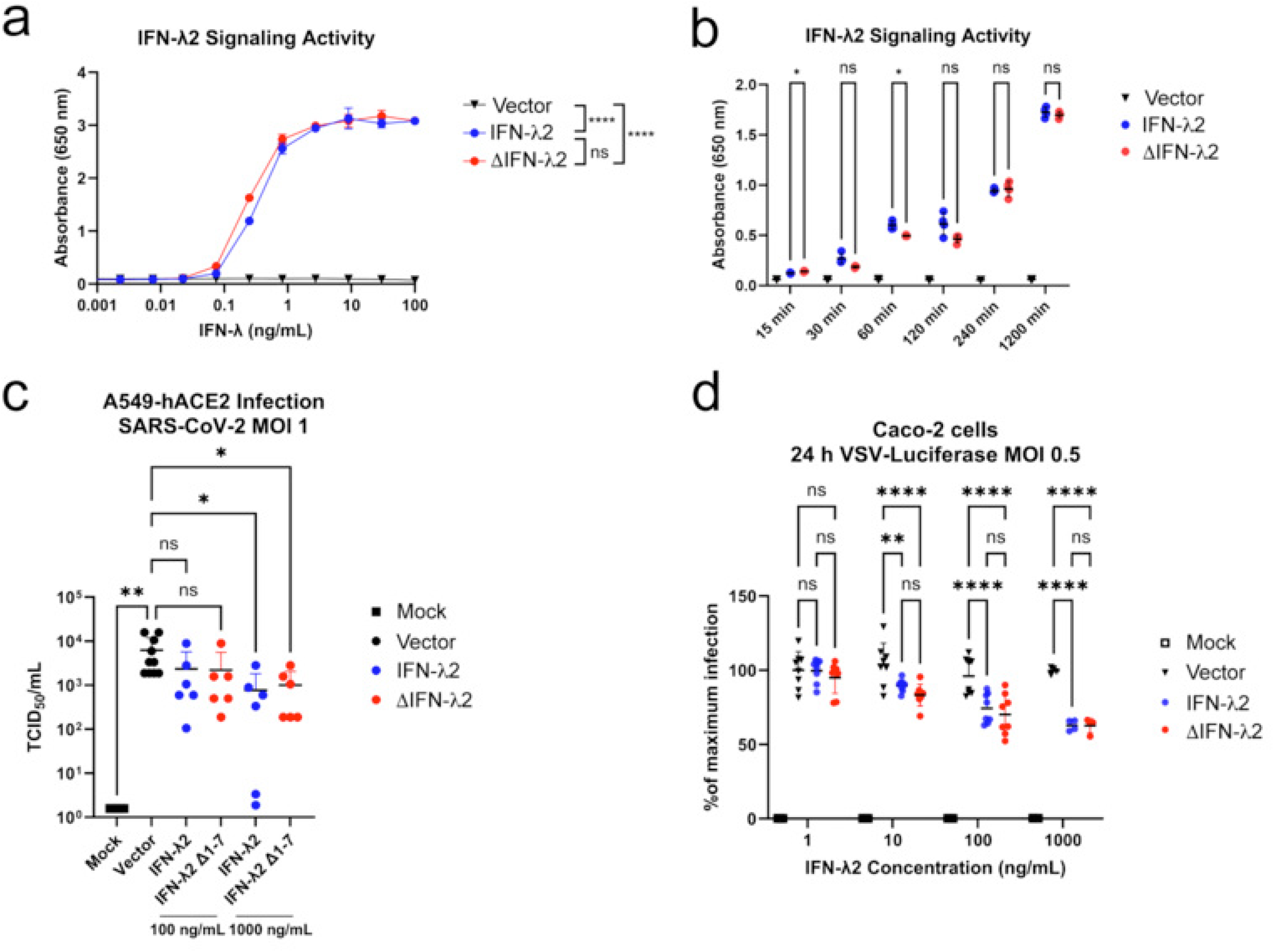
IFN-λ2 cleavage by 3CL^pro^ does not affect antiviral activity. (a and b) HEK-Blue IFN-λ signaling reporter cells were treated for (a) 24 h with increasing concentrations (*n* = 2 per group) or (b) over time with 0.3 ng/mL of IFN-λ2 or 3CL^pro^-cleaved IFN-λ2 analogue (ΔIFN-λ2) (*n* = 4). (c) A549 cells expressing human angiotensin-converting enzyme 2 (ACE2) were treated for 24 h with 100 or 1000 ng/mL of IFN-λ2, ΔIFN-λ2, or vector control, then infected with SARS-CoV-2 at a MOI of 1 for 48 h (*n* = 6 – 10, *N* = 2 independent experiments). Viral replication titre is reported as 50% tissue culture infectious dose per mL (TCID_50_/mL). (d) Caco-2 cells were treated for 24 h with 0 – 1000 ng/mL of IFN-λ2 or ΔIFN-λ2, then infected with replication-deficient (ΔG) vesicular stomatitis virus expressing firefly luciferase (VSV-luciferase) at a multiplicity of infection (MOI) of 0.5 for 24 h to assess antiviral activity (*n* = 4 – 8, *N* = 2). Statistical analysis was performed using (a, b, and d) two-way or (c) one-way ANOVA with Dunnett’s or Tukey’s post-hoc tests (*p < 0.05, **p < 0.01, ****p < 0.0001).

To further assess net antiviral activity after cleavage of IFN-λ2, A549 cells expressing the SARS-CoV-2 receptor human angiotensin converting enzyme 2 (hACE2) were pre-treated with IFN-λ2 or ΔIFN-λ2 for 24 h and then infected for 48 h with SARS-CoV-2 at a MOI of 1. Both IFN-λ2 and ΔIFN-λ2 similarly decreased SARS-CoV-2 viral titres, quantified as 50% tissue culture infectious dose (TCID_50_)/mL reflecting similar antiviral activity (Fig. 3c). A high throughput vesicular stomatitis virus (VSV) infection assay with luciferase reporter (53) again confirmed that both IFN-λ2 and ΔIFN-λ2 dose-dependently and similarly decreased VSV infection in Caco-2 cells across a broad range of concentrations from 1 – 1000 ng/mL (Fig. 3d). Taken together, 3CL^pro^ cleavage and removal of the N-terminal 7 residues of IFN-λ2 does not inhibit its IFN-λ signaling driven by the ISRE promoter or antiviral activity in two virus infection models.

### 3CLpro cleavage of IFN-**λ**1 blunts ISG protein expression responses

To evaluate the effects of 3CL^pro^ cleavage of IFN-λ1 on IFN receptor signaling, we treated Calu-3 (lung adenocarcinoma) cells and HepG2 (hepatocellular carcinoma) cells with IFN-λ1 or 3CL^pro^-cleaved ΔIFN-λ1 (ΔIFN-λ1) and observed decreased STAT1 phosphorylation of Tyr^701^ after cleavage (Fig. 4a). STAT1 phosphorylation was similarly decreased by increasing Ruxolitinib concentrations in Caco-2 and Calu-3 cells irrespective of 3CL^pro^ cleavage, confirming JAK-STAT signaling (Fig. 4b). Furthermore, intact IFN-λ1 induced the specific ISGs MX1, OAS2, and IFIT1 after 24 h in Calu-3 cells, whereas 3CL^pro^ cleavage abolished these ISG responses (Fig. 4c). Other ISGs, including iNOS, RSAD2, BST2, and SOCS1, showed no change in protein levels reflecting the specificity of IFN-λ signaling (Fig. S3). Therefore, SARS-CoV-2 3CL^pro^ cleavage of IFN-λ1 abrogates JAK1/2-mediated phosphorylation of STAT1 and specific ISG induction reproducibly across lung-, intestinal-, and liver-derived cell lines.

**FIG 4.**
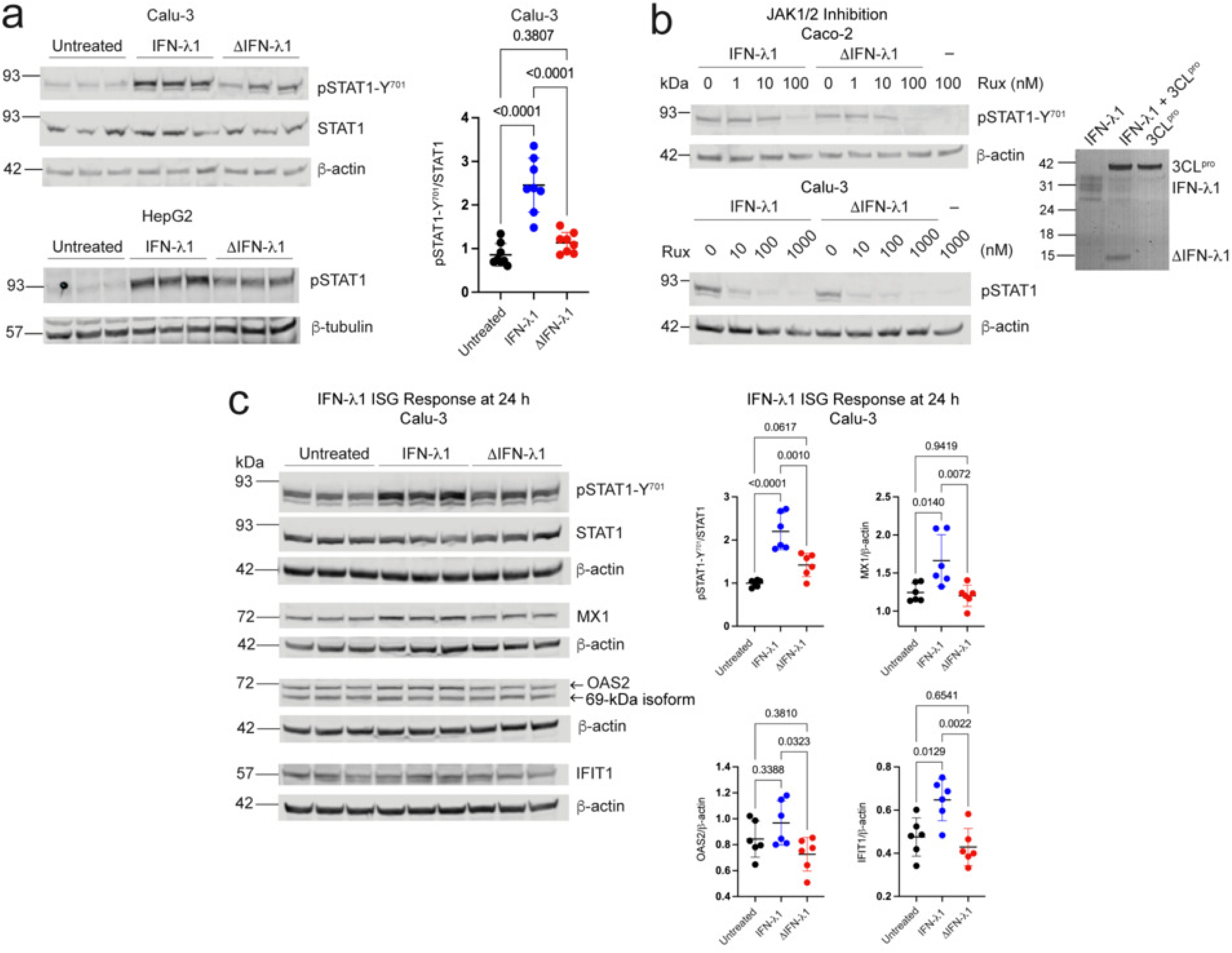
3CL^pro^ cleavage of IFN-λ1 dampens STAT1 phosphorylation, JAK-STAT signaling, and the ISG response. (a) Calu-3 lung epithelial cells (*n* = 8, *N* = 3) or HepG2 liver cells (*n* = 3) were treated with 1 µg/mL intact IFN-λ1 or 3CL^pro^-cleaved IFN-λ1 (ΔIFN-λ1) for 1 h at 37°C and lysates were immunoblotted for pSTAT1-Y^701^, STAT1, β-tubulin or β-actin. Densitometric quantification of pSTAT1-Y^701^ relative to STAT1 in IFN-λ1 treated Calu-3 cells (*n* = 8 per group, *N* = 3). Bars represent mean ± SD; statistical analysis using one-way ANOVA with Tukey’s post-hoc test and p < 0.05 significance threshold. (b) Caco-2 (top left panel) or Calu-3 (bottom left panel) cells were pre-treated for 2 h with increasing concentrations of JAK1/2 inhibitor Ruxolitinib (Rux), then treated for 1 h at 37°C with intact IFN-λ1 or ΔIFN-λ1, and lysates were immunoblotted for pSTAT1-Y^701^ and β-actin. Coomassie-stained 12% SDS-PAGE gel confirming IFN-λ1 cleavage by 3CL^pro^ prior to treatment of cells (right panel). (c) Calu-3 lung epithelial cells were treated for 24 h with intact or 3CL^pro^-cleaved ΔIFN-λ1 and lysates were immunoblotted for pSTAT1-Y^701^, STAT1, or for the ISGs MX1, OAS2, and IFIT1, with β-actin as a loading control on each blot (*n* = 6, *N* = 2). Densitometric quantification of pSTAT1-Y^701^ relative to STAT1, and ISGs relative to β-actin in Calu-3 lysates (*n* = 6, *N* = 2). Bars represent mean ± SD; statistical analysis using one-way ANOVA with Tukey’s post-hoc test and p < 0.05 significance threshold. Immunoblot antibodies: anti-pSTAT1-Y^701^ (1:1000), anti-β-actin (1:2000), anti-STAT1 (1:1000), anti-β-tubulin (1:2000), anti-MX1 (1:500), anti-OAS2 (1:500), anti-IFIT1 (1:500). See Fig. S8 for full uncropped gels and immunoblots.

### Glycosylation of IFN-**λ**1 is required for 3CL^pro^ cleavage

Since IFN-λ1 was not cleaved to completion, even with high concentrations of 3CL^pro^, and migrates as four separate bands on SDS-PAGE (Fig. 2a), we addressed if glycosylation of IFN-λ1 reduces cleavage by 3CL^pro^. Treatment of recombinant human IFN-λ1 with PNGase F to remove N-linked glycosylation, or with a combination of O-glycosidase and α2-3,6,8 neuraminidase to remove O-linked glycosylation, increased the electrophoretic migration of IFN-λ1 proteoforms on 12% SDS-PAGE (Fig. 5a), confirming that IFN-λ1 is N- and O-glycosylated.

**FIG 5.**
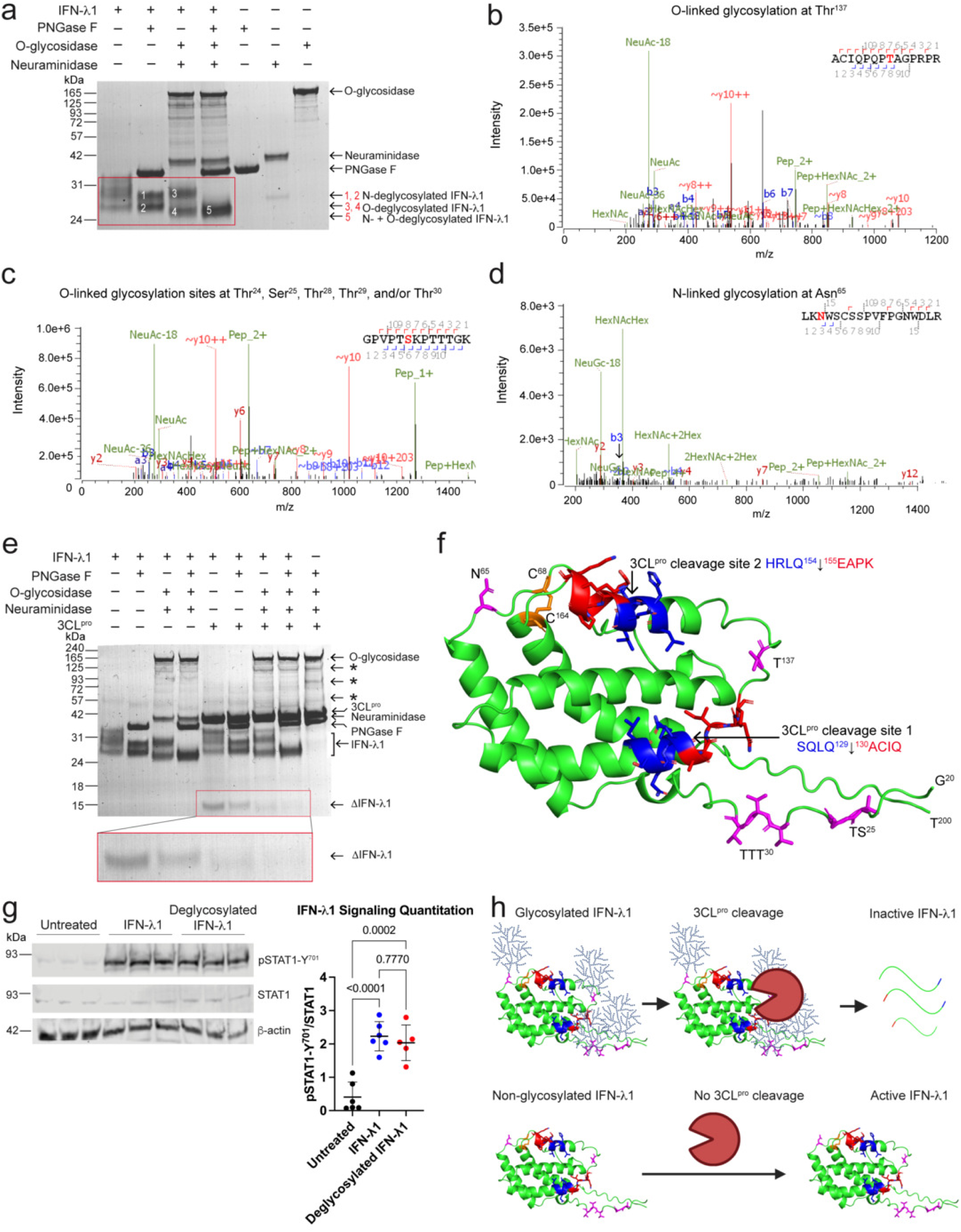
O-glycosylation of IFN-λ1 is required for 3CL^pro^ cleavage. (a) Coomassie-stained 12% SDS-PAGE gel of recombinant human IFN-λ1 treated for 20 h at 37°C with PNGase F to remove N-linked glycans, or O-glycosidase and neuraminidase to remove O-linked glycans, or all three deglycosylating enzymes (*N* = 2 independent experiments). The red box highlights IFN-λ1 under each deglycosylation condition compared with native IFN-λ1 control. (b–d) LC-MS/MS spectra showing (b) O-linked glycosylation at T^137^, (c) O-linked glycosylation site(s) at one or more Ser/Thr residues in the N-terminal region ^20^GPVPTSKPTTT, and (d) an N-linked glycosylation site at N^65^ with the arrow pointing to the m/z peak of the b3 ion. (e) 12% SDS-PAGE analysis of recombinant human IFN-λ1 incubated for 24 h at 37°C in buffer, or with PNGase F, O-glycosidase or neuraminidase, or all three deglycosylating enzymes, and then incubated for a further 20 h at 37°C with or without 5 µM 3CL^pro^ (*N* = 2). The red box highlights cleaved IFN-λ1 under each deglycosylation condition. (f) AlphaFold structure prediction of IFN-λ1 showing experimentally identified glycosylation sites (magenta), disulphide bond (orange), non-prime side of each 3CL^pro^ cleavage site (blue) and prime side of each 3CL^pro^ cleavage site (red) (model ID: AF-Q8IU54-F1). (g) Caco-2 epithelial cells were treated with intact or deglycosylated IFN-λ1 for 1 h at 37°C, and lysates were immunoblotted for pSTAT1-Y^701^, STAT1, and β-actin (*n* = 6, *N* = 2). Densitometric quantification of pSTAT1-Y^701^ relative to STAT1 in Caco-2 lysates (mean ± SD; *n* = 6, *N* = 2) with statistical significance assessed using one-way ANOVA with Tukey’s post-hoc test (p < 0.05 significance threshold). (h) Model showing that glycosylated IFN-λ1 is cleaved by 3CL^pro^ at the two cleavage sites nearby O-glycosylated Thr^137^ or N-terminal site(s), whereas non-glycosylated IFN-λ1 is resistant to cleavage by 3CL^pro^. See Fig. S9 for full uncropped gels and immunoblots.

LC-MS/MS analysis identified two O-glycosylation sites: one at Thr^137^ (Fig. 5b), and one near the flexible N-terminus of IFN-λ1 at one or more Ser or Thr residues in the sequence ^20^GPVPTSKPTTTGK^32^ (Fig. 5c, S4). We also found peptide spectral matches confirming N-linked glycosylation at Asn^65^ (Fig. 5d). As the O-linked glycosylated chains are near cleavage sites 1 and 2, we assessed whether glycosylation may block access and cleavage of IFN-λ1 by 3CL^pro^. We first incubated IFN-λ1 for 24 h in combinations or absence of PNGase F, O-glycosidase, and α2-3,6,8 neuraminidase, and then treated the deglycosylated IFN-λ1 for 20 h with 3CL^pro^ or buffer. Unexpectedly, rather than retarding cleavage, O-glycosylation was essential for efficient cleavage of IFN-λ1 by 3CL^pro^ (Fig. 5e). N-glycosylation still enhanced cleavage but had less of an effect than the O-linked chains that are positioned proximal to cleavage site 1 (Fig. 5f). These results suggest a model that binding of 3CL^pro^ to O-linked glycan chains either near the N-terminus of IFN-λ1 or at Thr^137^, facilitates cleavage.

To determine the effect of glycosylation on IFN-λ1 signaling through the JAK-STAT pathway, we treated Caco-2 cells with native or deglycosylated IFN-λ1 and found no significant differences in pSTAT1 signaling (Fig. 5g). Whereas glycosylation is not essential for IFN-λ1 signaling, it does tether and enhance cleavage of IFN-λ1 by 3CL^pro^. Indeed, the absence of glycosylation protects that fraction of the IFN-λ1 pool from proteolysis by SARS-CoV-2 3CL^pro^ (Fig. 5h).

### MMPs cleave 3CL^pro^, but 3CL^pro^ does not cleave MMPs

Several MMPs, particularly MMP2, MMP7, MMP8, and MMP9, have been identified as biomarkers of COVID-19 severity (47, 58–61). We investigated potential cross-inactivation mechanisms between MMPs and 3CL^pro^. To test this, we incubated inactive 3CL^pro^ (Cys145Ala) as a substrate with human MMP1, MMP2, MMP7, MMP8, MMP9, MMP12 or MMP14 and used SDS-PAGE to resolve cleavage fragments. In particular, MMP7 and MMP12, which are produced by monocytes and macrophages, respectively (62, 63), were the most effective (Fig. 6a), followed by MMP2 and MMP8. MMP2 is expressed by many cells including endothelial cells and fibroblasts, whereas MMP8 is essentially a neutrophil-specific protease (64, 65). MMP1 and MMP14 only poorly cleaved 3CL^pro^, whereas MMP9 did not cleave 3CL^pro^ (Fig. 6a). In the converse experiment, we inhibited the MMPs by the broad-spectrum MMP inhibitor marimastat at >10-fold molar excess as substrate for 3CL^pro^ over 18-h incubation. 3CL^pro^ did not cleave any of the MMPs (Fig. 6b, c), and was not inhibited by marimastat (Fig. S5). Together, these results suggest a model whereby MMPs may function to counteract extracellular 3CL^pro^ by proteolytic degradation (Fig. 6d).

**FIG 6.**
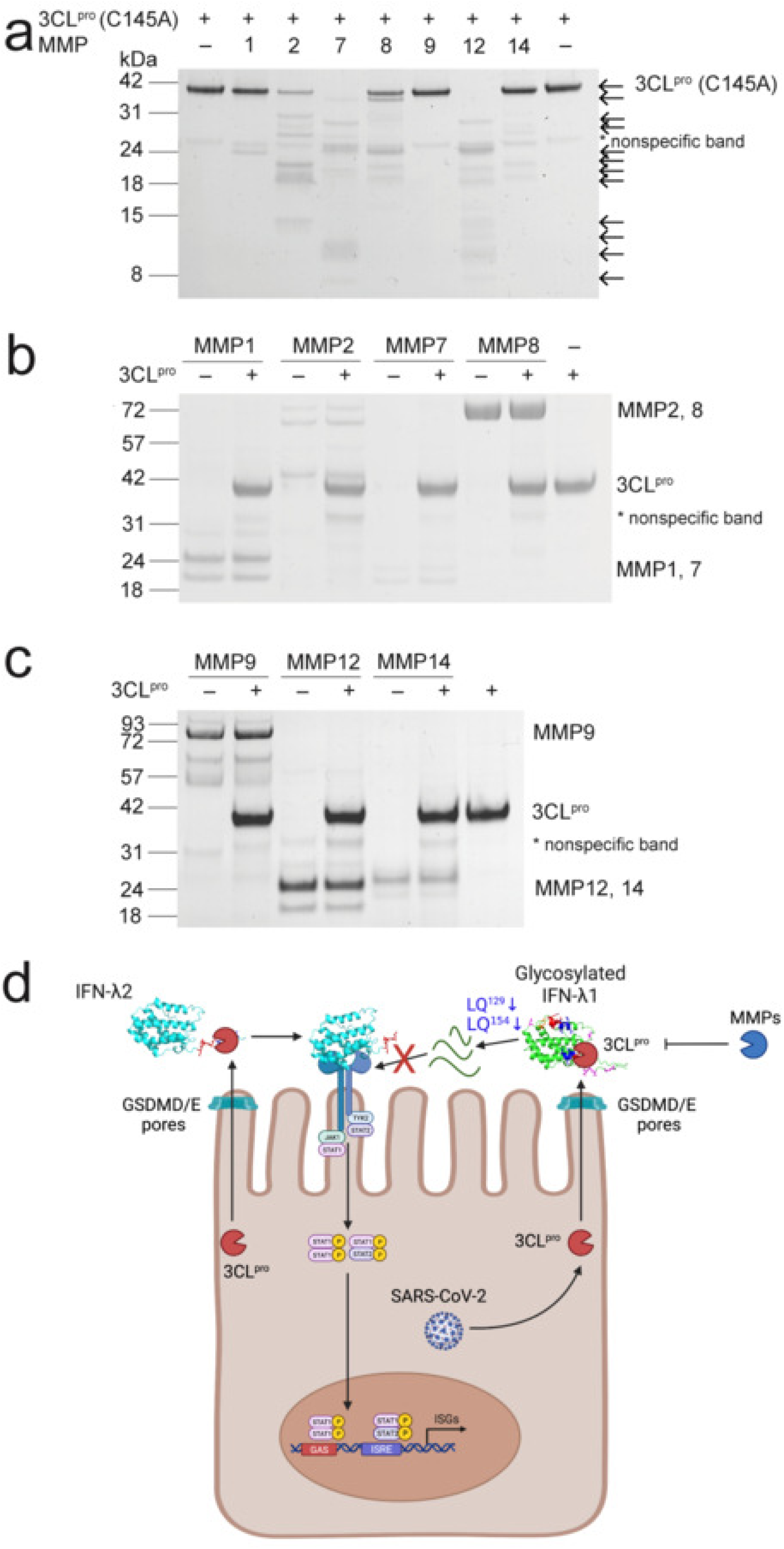
Matrix metalloproteinases (MMPs) cleave 3CL^pro^, but 3CL^pro^ does not cleave MMPs. (a) Coomassie-stained 12% SDS-PAGE gel of 3 µM 3CL^pro^ (Cys145Ala) inactive mutant after 20 h incubation alone or with 300 nM of MMP1, MMP2, MMP7, MMP8, MMP9, MMP12, or MMP14 at 37°C (representative of *N* = 3 independent experiments). Arrows show cleavage fragments produced by each MMP. (b) Coomassie-stained 12% SDS-PAGE gel showing 2 µg of MMP1, MMP2, MMP7, or MMP8 incubated with MMP-inhibitor marimastat (50 µM) and with or without 3CL^pro^ (2.5 µM) for 20 h at 37°C. (c) Coomassie-stained 12% SDS-PAGE gel of MMP9, MMP12, or MMP14 (2 µg) incubated with MMP-inhibitor marimastat (50 µM) and with or without 3CL^pro^ (2.5 µM) for 20 h at 37°C. (d) Model of 3CL^pro^ secretion from SARS-CoV-2-infected host cells through gasdermin-D/E (GSDMD/E) pores. Extracellularly, 3CL^pro^ cleaves IFN-λ2 at LH^32^↓^33^G without altering its antiviral activity and inactivates O-glycosylated IFN-λ1 by proteolysis. Extracellular host MMPs, such as MMP2, MMP7, MMP8, and MMP12 cleave and inactivate 3CL^pro^, thereby blocking IFN-λ1/2 cleavage. See Fig. S10 for full uncropped gels.

## DISCUSSION

Our results identifying SARS-CoV-2 3CL^pro^ cleavage of IFN-λ1 and IFN-λ2 type III interferons increase the breadth of proteolytic regulation of interferons that has been previously reported for type I and type II interferons by MMP12 and MMP9, respectively (44–46). The irreversible inactivation of interferons adds a further layer of complexity in the regulation and antagonism of these antiviral cytokines. In extracellular tissue microenvironments of the nasopharynx, upper respiratory tract, or gastrointestinal tract at sites of SARS-CoV-2 infection, cleavage of IFN-λ1 by 3CL^pro^ may contribute to suppression of the immune response by SARS-CoV-2, potentially compromising epithelial barrier immunity and promoting the spread of infection. Our findings that O-glycosylated IFN-λ1 was inactivated by 3CL^pro^, IFN-λ2 was cleaved by 3CL^pro^ yet remained active, and IFN-λ3 and IFN-λ4 were resistant to 3CL^pro^ cleavage together suggest that evolutionary redundancy in type III IFN signaling in humans via multiple IFN-λs may aid in maintaining epithelial barrier defences during infections despite extracellular viral protease activity. The inactivation of O-glycosylated IFN-λ1 signaling by 3CL^pro^ cleavage may contribute to the delayed IFN-λ1 response observed during SARS-CoV-2 infection (66, 67).

MMPs are critical regulators of inflammation by processing chemokines and cytokines (40–46, 68) and in turn, our present results show that inflammatory and tissue MMPs degrade 3CL^pro^, revealing their potential to antagonize the activity of extracellular 3CL^pro^ in some settings. Interestingly, the tissue inhibitor of metalloproteinases (TIMP)-1 levels have been positively correlated with COVID-19 disease severity in clinical studies (69, 70). Moreover, several MMPs, particularly MMP2, MMP7, MMP8, and MMP9, have been identified as biomarkers of COVID-19 severity (47, 58–61).

We show here that a 6-residue C-terminal peptide is removed by 3CL^pro^ cleavage of IFN- γ (Arg160Gln). IFN-γ signaling through STAT1 phosphorylation was unchanged following cleavage, which is interesting because cleavage of the C-terminus at E^135^↓L and M^157^↓L by MMP12 (44) and the C-terminal 11 residues by bacterial proteases (71) inhibits IFN-γ binding to its receptor, evidencing the importance of residues 156 to 160 for IFN-γ signaling. Recent structures of the heterohexameric IFN-γ:IFN-γ:IFNγR1:IFNγR2 signaling complex (72) did not resolve this C-terminal disordered region of IFN-γ, which likely undergoes a key binding interaction necessary for downstream signaling.

Specifically, we identified 3CL^pro^ cleavage sites in the N-terminus of IFN-λ2 at LH^32^↓G, in the C-terminus of IFN-γ (Arg160Gln) at FQ^160^↓G, and two sequential cleavages in IFN-λ1 at LQ^129^↓A then at LQ^154^↓E resulting in loss of activity. This is a new mechanism to antagonize host antiviral mechanisms that other viral proteases expressed by *Picornaviridae, Flaviviridae, Adenoviridae,* and *Poxviridae* among other virus families may also employ after protease secretion through gasdermin D or gasdermin E pores.

We further show that IFN-λ1 is O-glycosylated at Thr^137^ and at one of several Ser or Thr residues in the sequence ^24^TSKPTTT^30^, and is N-glycosylated at Asn^65^. The variety of glycosylated proteoforms of IFN-λ1 may regulate half-life, bioavailability, metabolism, or clearance *in vivo.* We found that glycosylation of IFN-λ1 is important because 3CL^pro^ efficiently cleaved only O-glycosylated IFN-λ1 whereas non-glycosylated IFN-λ1 was resistant to cleavage. This was unexpected. Glycosylation improves the stability and activity of proteins from proteolytic degradation *in vivo*, including type I IFNs (73, 74), and in lieu of glycosylation, recombinant proteins are often pegylated to provide similar protection (75). Nonetheless, isolated examples of enhanced cleavage of substrates has been reported. MMPs were shown to bind to the O-glyco-side chain conjugated to non-prime side P1 residue in substrates that enhanced cleavage (76). Tethering of proteases at or near their cleavage site motifs increase their specific activity through localised increased concentration resulting in zero order kinetic activity.

Loss of O-glycosylation led to a general decrease in proteolysis by human MMPs (76), which is consistent with our finding that O-glycosylation of IFN-λ1 is important for cleavage by 3CL^pro^. Substrate-binding exosites on 3CL^pro^ may recognize the N-terminal O-linked glycosylation unstructured ^24^TSKPTTT^30^, to bring 3CL^pro^ into a more favourable position to cleave IFN-λ1. Alternatively, binding to O-glycosylated Thr^137^ may facilitate cleavage of the scissile bond located 8 residues away in cleavage site 1 through increased local concentration effects. These observations suggest that an outcome of variable glycosylation of IFN-λ1 may be to limit susceptibility to protease-mediated inactivation while retaining critical signaling functions. In conclusion, our study highlights the robustness of the antiviral IFN system through gene and protein isoform redundancy including variable glycosylation of IFN-λ1 to limit inactivation by viral proteases.

## ACKNOWLEDGMENTS

We thank Michael Berne, Tufts University School of Medicine, for performing N-terminal sequencing using Edman degradation, and Yoan Machado and Peter Bell, University of British Columbia, for assistance with experimental design, substrate cleavage site prediction and peptide design for protease cleavage assays, and MALDI-TOF. We thank Gert Zimmer at the Institute for Virology and Immunology for gifting luciferase-expressing VSV, and Caroline Lehmann for technical support with IFN-λ2 full-length and ΔIFN-λ2 expression and quantification. This work was supported by the Canada Research Chairs program (950-01-126 to CMO), the Canadian Institutes for Health Research (CIHR) Foundation Grant program (FDN-148408 to CMO), the C.I.H.R. Canadian 2019 Novel Coronavirus (COVID-19) Rapid Research Funding Opportunity (F20-01013 to Dr. Eric Jan, University of British Columbia and CMO), the Canada Foundation for Innovation (31059 to CMO), the Michael Smith Foundation for Health Research (IN-NPG-00105 to CMO), CIHR Banting and Best Canada Graduate Doctoral Award (201911 FBD-434976-268115 to PMG), the Centre for Blood Research (Graduate Student Award to PMG), the University of British Columbia (Four-Year Fellowship to PMG) and the Swiss National Science Foundation (310030-212380 to CB).

## AUTHOR CONTRIBUTIONS

PMG designed and performed the experiments, analyzed data, prepared figures, wrote and revised the manuscript; HCR performed LC-MS/MS analyses; IP designed the recombinant 3CL^pro^ purification protocol and purified recombinant 3CL^pro^; RK purified recombinant 3CL^pro^ and 3CL^pro^ (Cys145Ala), ordered reagents, and managed the laboratory; GSB purified recombinant 3CL^pro^ (Cys145Ala), contributed to experimental design, and revised the manuscript; CB designed experiments and interpreted data, obtained funding, and revised figures and the manuscript; CMO conceived the study, designed experiments and interpreted data, obtained funding and supervised the study, revised figures and the manuscript. All authors approved the final version for publication. All authors contributed in a meaningful manner to the research presented in this manuscript. All authors have given approval to the final version of the manuscript. The author’s order was determined on the basis of the quantity of research and data analysis and preparation performed.

## Notes

The authors declare no competing financial interest.

## FUNDING

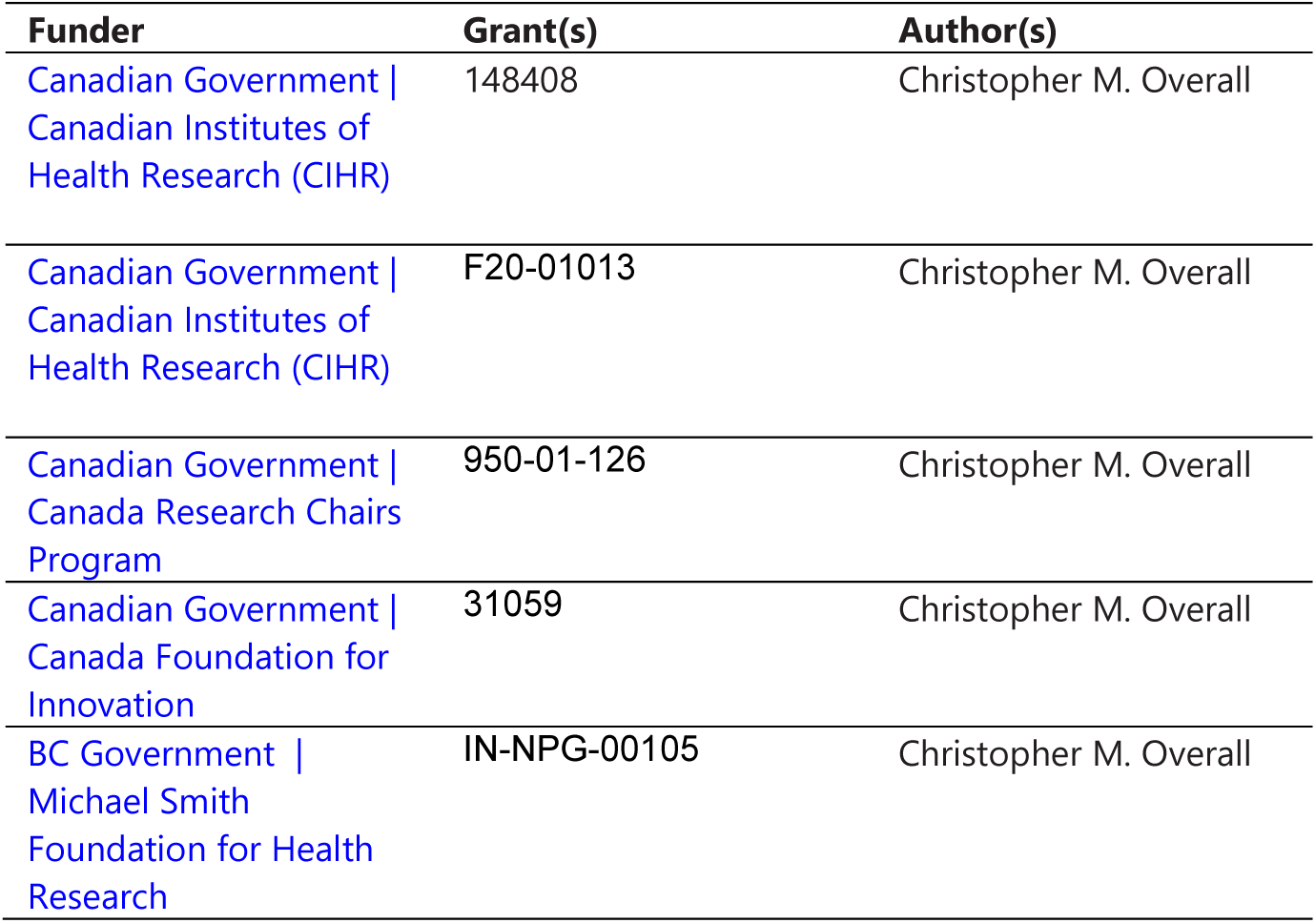
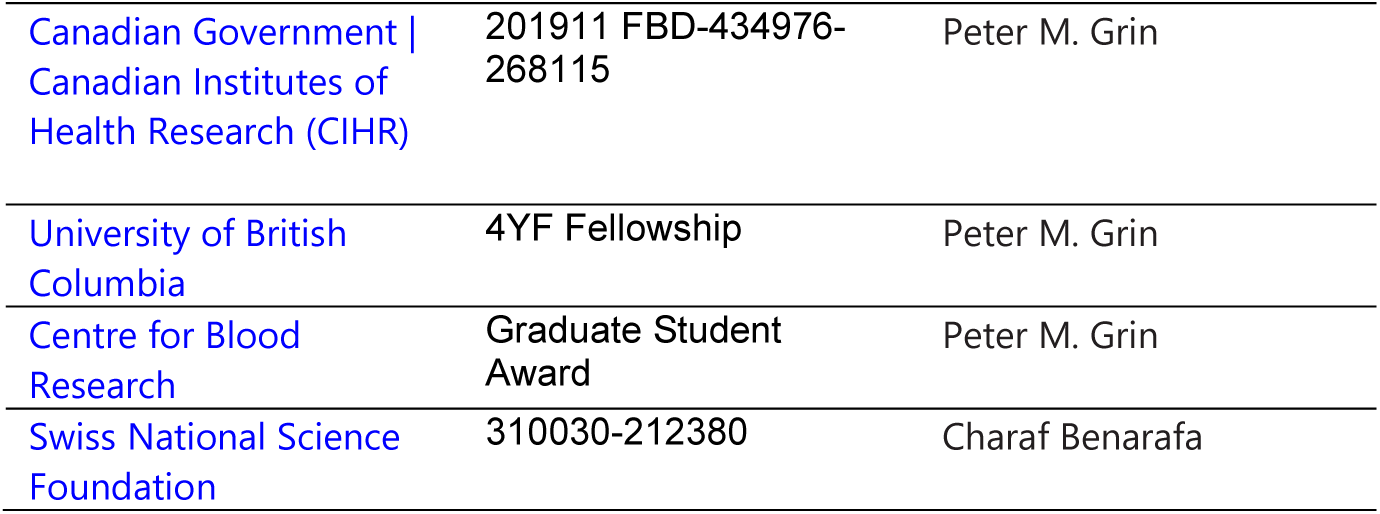

## DATA AVAILABILITY

Further information and requests for resources and reagents can be directed to Professor Chris Overall.

## Supplemental Material

Figures S1 to S10.

